# Conserved Pseudouridines in Helix 69 of the Ribosome are Important for Ribosome Dynamics in Translation

**DOI:** 10.1101/2024.02.02.578723

**Authors:** Hong Jin, Xin Chen

**Affiliations:** Department of Biochemistry, University of Illinois at Urbana-Champaign, 600 S. Mathews Avenue, Urbana, IL 61801; Center for Biophysics and Quantitative Biology, University of Illinois at Urbana-Champaign, 600 S. Mathews Avenue, Urbana, IL 61801; Carl R. Woese Institute for Genomic Biology, 1206 West Gregory Drive, University of Illinois at Urbana-Champaign, 600 S. Mathews Avenue, Urbana, IL 61801

**Author notes:** Equal contributions.

**Keywords:** Translation, Ribosome, Ribosome dynamics, Helix 69, RNA modification, Pseudouridine, Pseudouridylation, snoRNA

## Abstract

The widespread distribution of pseudouridine (Ψ), an isomer of the canonical uridine base, in RNA indicates its functional importance to the cell. In eukaryotes, it is estimated that around 2% of ribosomal RNA nucleotides are pseudouridines, most of which are located in functional regions of the ribosome. Defects in RNA pseudouridylation induce a range of detrimental effects from compromised cellular protein biosynthesis to disease phenotypes in humans. However, genome-wide changes to mRNA translation profiles by ribosomes lacking specific conserved pseudouridines have not been extensively studied. Here, using a new genomic method called 5P-Seq and *in vitro* biochemistry, we investigated changes in ribosome dynamics and cellular translation profiles upon loss of Ψ2258 and Ψ2260 in helix 69, the two most conserved pseudouridines in the ribosomes. We found that inhibiting the formation of these two pseudouridines challenges ribosomes to maintain the correct open reading frame and causes generally faster ribosome dynamics in translation. Furthermore, mutant ribosomes are more prone to pause while translating a subset of GC-rich codons, especially rare codons such as Arg (CGA) and Arg (CGG). Our data suggest the presence of Ψ2258 and Ψ2260 contributes to the dynamics of the H69 RNA stem-loop, and helps to maintain functional interactions with the tRNAs as they move within the ribosome. The optimality of this ribosome-tRNA interaction is likely to be more critical for those limited tRNAs that decode rare codons. Consistent with the changes in ribosome dynamics, our data also show that IRES-mediated translation is compromised in the mutant ribosome. These results explain the importance of Ψ2258 and Ψ2260 in H69 to maintain cellular fitness. The strong conservation of Ψ2258 and Ψ2260 in the ribosomes from bacteria to humans indicates their functional significance in modulating ribosome functions. We anticipate that the identified functions of these covalent modifications will be conserved in other species.

## INTRODUCTION

Proteins in all living cells are synthesized in the remarkable cellular machine called ribosome. Beyond their traditional role as the protein synthesis factory, ribosomes integrate different signals and communicate with other pathways to regulate gene expression (1-3). Despite variations in size and composition, all ribosomes across different species consist of two asymmetrical subunits that carry out different, but closely related functions in translation. The small subunit (30S in bacteria; 40S in eukaryotes) decodes the genetic information in the messenger RNA (mRNA), and the large subunit (50S in bacteria; 60S in eukaryotes) makes peptide bonds in the protein. Interactions between the two ribosomal subunits occur via so-called “bridges”, which were first characterized by cryo-electron microscopy (cryo-EM)(4) and subsequently characterized in molecular detail by crystallography (5, 6). Inter-subunit bridges of the ribosome not only “glue” the two subunits together, they also closely interact with tRNAs, translation factors, and other ligands. Furthermore, these bridges critically control the ratcheting movement that is characterized by relative subunit rotation, a large-scale conformational re-arrangement in the ribosome that is fundamentally important for ribosomal function in each step of translation and its regulation.

Many bridge contacts in the ribosome, especially those that are centrally located, are mediated by highly conserved RNA-RNA interactions. Taking the essential B2a bridge near the decoding center of the small ribosomal subunit as an example, this bridge is formed by the conserved interaction between the tip of Helix 69 (H69, **Figure 1A**) of the 28S rRNA and the minor groove of helix 44 (h44) of the18S rRNA. From bacteria to humans, when the small subunit rotates relative to the large one during the ratcheting movement of the ribosome (7, 8), the B2a bridge contact is maintained by an approximately 6 Å compression of H69 to accommodate the shift of h44 toward the E site. This bridge is important for translocation and translation fidelity, as it contacts the D stem region of the A- and P-site tRNAs (7-9). Nucleotide mutations in H69, even a point mutation in H69 loop region (U1915A), caused strong frameshifts and stop-codon read-through on reporter RNAs (10, 11).

**Figure 1.**
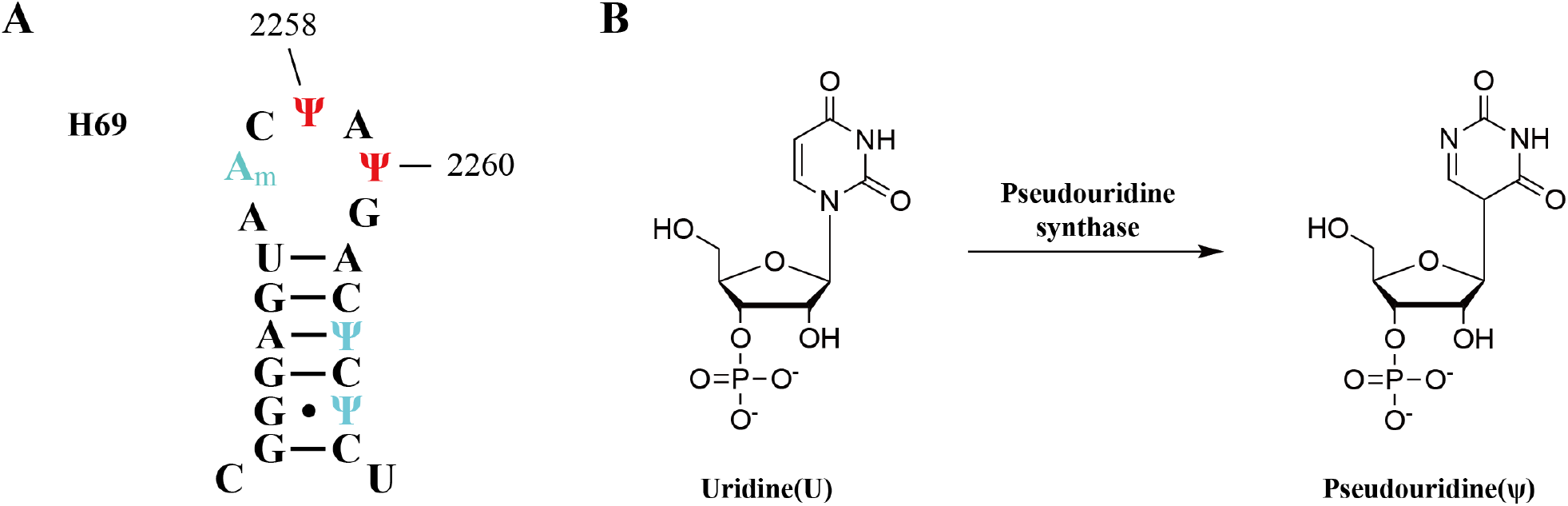
Pseudouridines and RNA pseudouridylation. A. **Nucleotide sequence and secondary structure of H69 of *S. cerevisiae* ribosome.** Modified nucleotides in H69 are colored. Two most conserved ribosomal Ψ2258 and Ψ2260 of interest in this study are indicated in red. B. **RNA pseudouridylation.** Conversion of uridine to pseudouridine by pseudouridine synthase in the cell.

H69 is rich in RNA modifications. In fact, the primary transcript of ribosomal RNA (pre-rRNA) undergoes extensive, site-specific covalent modification, mainly 2’-O-ribose-methylations and replacement of uridines by pseudouridines (**Figure 1B**) (12). rRNA modifications have long been of great interest because of their abundance and conservation. Importantly, most of the modifications are clustered in the functionally important regions of the ribosome. Variations of rRNA modifications including defects in rRNA methylation (13, 14) and pseudouridylation (15, 16) were shown to be detrimental to the cell, but molecular details accounting for these phenomena remain to be better understood. Loss of one or multiple rRNA modifications in or near the functionally important regions of the ribosome was reported to impair cell growth, mature ribosome formation, and affect translation accuracy on specific reporters (17-20). However, changes in the translation profiles of cellular mRNAs at a genome-wide scale by mutant ribosomes have not been extensively studied.

In this study, using a genomic approach called 5P-Seq to examine *in vivo* ribosome dynamics, we report changes in mRNA translation profiles when ribosomes lose two conserved pseudouridines, Ψ2258 and Ψ2260, in H69. These two Ψs are among the most conserved rRNA modifications in the ribosome from bacteria to humans. However, the deletion of these two modifications does not affect the viability of the cell under standard laboratory conditions, although the mutant cell cannot compete with the wild type (18, 21). Here, we demonstrate that inhibiting pseudouridinylation at these two conserved positions results in faster overall ribosome dynamics, renders the translation machinery prone to frameshifting, and affects the movement of tRNAs decoding CG-rich codons in ribosomal A and P sites. Of note, the loss of these two conserved pseudouridines in an otherwise unaltered ribosome compromises the IRES-mediated translation. These results not only provide a more comprehensive view of the altered translation profile in mutant cells, but also provide an explanation as to how rRNA modifications contribute to cell fitness.

## RESULTS

In archaea and eukaryotes, rRNA modifications are catalyzed by small nucleolar ribonucleoprotein (snoRNP) complexes. The protein components of the snoRNP confer a generalized methylation and pseudouridylation activity to the complex. The RNA components, snoRNAs, define the sequence specificity of a snoRNP by base-pairing with the target RNA and directing the complexed protein nucleotide-modification machinery to the appropriate nucleotide for covalent modification. snoRNAs are classified into two major families based on their secondary structure features: box H/ACA snoRNAs and box C/D snoRNAs, which guide 2’-O-ribose-methylation and pseudouridylation, respectively. A box H/ACA snoRNA, snR191, which is conserved from yeast to humans, is responsible for the formation of Ψ2258 and Ψ2260 in the ribosome (21).

To study the function of these two conserved pseudouridines (Ψs) in the ribosome and to gain insight into how cell fitness is compromised in their absence, snR191 was deleted (21) and the resulting mutant Saccharomyces cerevisiae will have ribosomes containing uridines instead of Ψs at positions 2258 and 2260 in H69 (**Figure 1A**). The mutant was named mH69 in this study. *in vivo* ribosome dynamics on mRNAs in WT and mH69 were compared using a new genomic method called 5P-Seq (22, 23). In this method, 5’ phosphorylated mRNA intermediates that are generated in the co-translational degradation process are sequenced. The position and the frequency of these intermediates infer ribosome dynamics in mRNA translation at a single-codon resolution (**Figure 2A**). Since 5P-Seq does not require the use of translation inhibitors or *in vitro* mRNA digestion, it offers an accurate depiction of ribosome footprints on translating mRNAs. We performed multiple independent 5P-Seq experiments with both mH69 (ie, ΔsnR191) and its isogenic WT strain, data quality and reproducibility are shown in **Supplemental Figure S1**.

**Figure 2.**
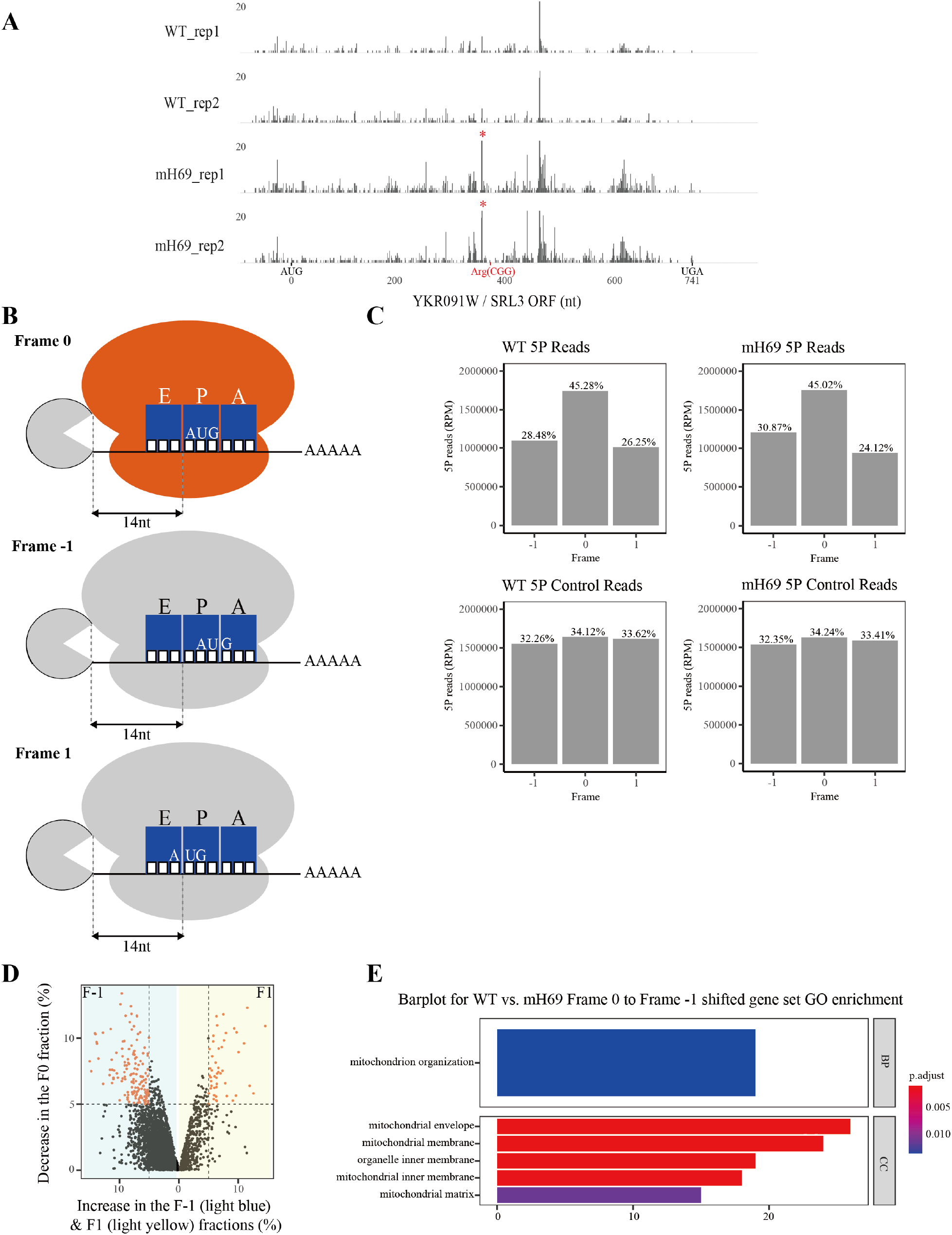
Loss of Ψ2258 and Ψ2260 in H69 challenges the ribosome to maintain correct open reading frames. **A Loss of Ψ2258 and Ψ2260 in H69 exacerbates ribosome stalling at rare codons**. Representative genome tracks of 5’ ends of 5P reads for WT and mH69 samples. The presence of Arginine (Arg) codon CGG within the YKR091W/SRL3 Open Reading Frame (ORF) is marked in red. Notably, a distinct stalling pattern is observed in the mH69 sample occurring 14 nt upstream of the Arg (CGG) site (Red *). **B. Illustration of ribosome-protected frames in the 5P-Seq experiment**. Schematic representation of three possible ribosome-protected frames in the 5P-Seq experiment. When the ORF of each gene is aligned with the ribosomal A-site and P-site, ie, 17 nt and 14 nt downstream of the 5’ end of reads of each gene are aligned to the ribosomal A-site and P-site, respectively, the ribosome-protected ORF frame is defined as Frame 0. One nucleotide frameshifting, upstream or downstream, from the correct ribosomal ORF is defined as Frame -1 and Frame 1, respectively, in this study. **C. Frame distribution of the 5P-Seq data in mH69 and WT**. Histograms representing the frame distribution of 5P reads in the coding regions of all mRNAs. Consistent with the ribosome-protected mRNA region during translation, 5P-Seq data are predominantly enriched in Frame 0. In contrast, control samples exhibit no distinct preference among F-1, F0 and F+1 frames, indicating random distribution of reads on mRNAs in the absence of active translation. **D. Loss of Ψ2258 and Ψ2260 of H69 challenges the ribosome to maintain the correct open reading frames**. Genes whose translation by ribosomes lacking Ψ2258 and Ψ2260 exhibit frameshifting are shown. These genes are defined as those with a decreased in F0 fraction (ie, the reads ratio of (F0/F(−1) + F0 +F(+1))) of 5% or larger. The frames that deviated from F0, either Frame 1 (F1) or Frame -1 (F-1), are determined by the more significantly increased reads in the corresponding frames. The y-axis denotes the decrease in the F0 fraction as the result of loss of Ψ2258 and Ψ2260. The x-axis denotes the increased fractions of -1 frameshifts (Light blue area) and +1 frameshifts (light yellow). A change of 5% or higher is considered to be significant in the data analysis, and these genes are shown in red. **E. Gene Ontology (GO) enrichment analysis for transcripts undergoing -1 frameshifting** The bar plot illustrates GO enrichment pathways for 152 genes that exhibit a significant shift from Frame 0 to Frame -1. The color gradient from red to purple represents an adjusted p-value ranging from low to high. The GO terms are listed on the y-axis, while the x-axis presents the number of genes associated with each given GO term. BP and CC represent Biological Process and Cellular Component, respectively. The GO terms with an adjusted p-value less than 0.05 are considered statistically significant.

### 1. Loss of Ψ2258 and Ψ2260 in the ribosome increases frameshifts during translation

In addition to the codon-specific ribosome protection pattern given by appropriate translation of a CDS from the start codon, reads generated by 5P-Seq also include those from the +1 and -1 frames when mRNA transcripts are degraded by the 5’ exonuclease (22). During the process of data analysis, we denote the sequenced reads with either the 14nt or 17nt position mapped to the ribosome-protected frame (ie, the ORFs starting at the start codon AUG) as Frame 0, and other reads that had deviated from Frame 0 as Frame -1 and +1 (**Figure 2B**). Compared to wild type (WT), mH69 shows an overall -1 frameshift, as indicated by a decrease in the reads covered in the correct ribosome-protected frame (Frame 0), and an increase in the number of reads detected in the -1 Frame (**Figure 2C**). This observation suggests that ribosomes missing Ψ2258 and Ψ2260 of H69 struggle to maintain the correct open reading frame. Among the significant variations (defined as >5% decrease in the F0 fraction at the given gene in mutants), we found that 273 mRNA transcripts lost their correct translation frame, of which 152 transcripts show a significant -1 frameshift and 49 mRNAs show a significant +1 frameshift (**Figure 2D)** Results from the GO analysis indicate that the -1 frameshifted mRNAs are enriched in genes functionally related to mitochondrial organization and components (**Figure 2E**). By contrast, transcripts that underwent the +1 frameshift are not particularly enriched in any particular groups or pathways.

### 2. Missing Ψ2258 and Ψ2260 results in faster translation initiation and termination

The clear 3-nt periodicity pattern in our mapped reads indicates the speed of ribosome movement on mRNAs. In the mapped data, an increased accumulation of 5P intermediates indicates slower ribosome movement, whereas a decreased number of 5P reads at the same position shows faster dynamics (22). Given the size of the exonuclease which contacts the last translating ribosome on an mRNA undergoing co-translational degradation, the observed peak at the -14 position (**Figure 3A**) corresponds to the pause of translation machinery undergoing start codon recognition during initiation. We noticed a decreased peak at -14nt in mH69 compared to the WT ribosome when we analyzed the 5P patterns surrounding the start codon (**Figure 3A**, the red peak at the -14 position). This observation suggests that at the metagene level, translation initiation is faster when Ψ2258 and Ψ2260 are absent in the ribosome.

**Figure 3.**
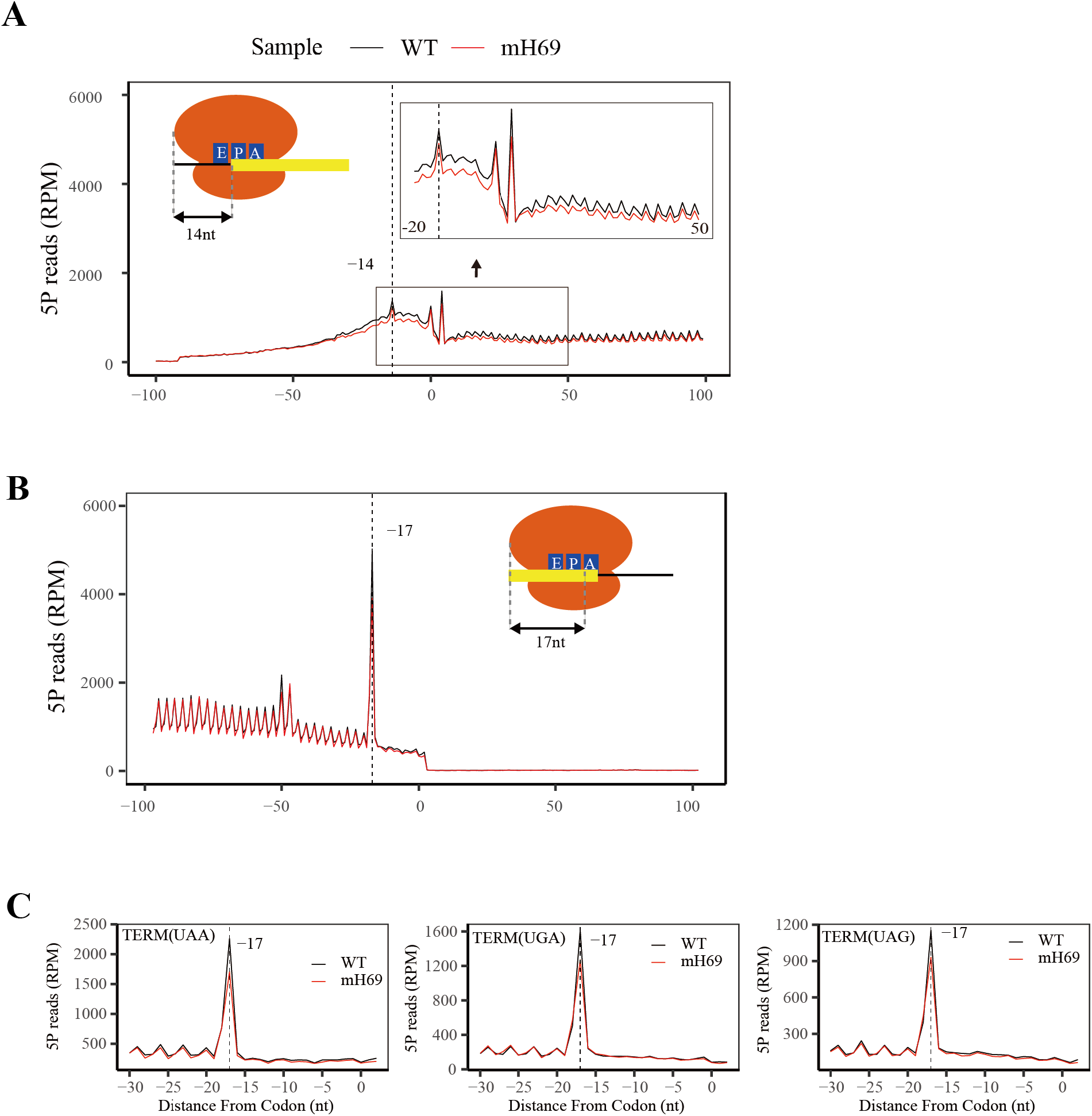
Inhibition of the formation of Ψ2258 and Ψ2260 in H69 results in faster ribosome dynamics in translation initiation and termination. **A. Metagene analysis displays the abundance of 5P reads relative to the start codon in ORF**. The metagene analysis illustrates the distribution of 5P reads around the start codon AUG region. The WT and mH69 are depicted in black and red, respectively. The peak at 14 nt upstream of the start codon in 5P-Seq experiment indicates the ribosome pause during initiation, after which the 3nt periodicity begins. The region extending from -20 nt to +50 nt relative to the start codon is enlarged in the insert. Of note, an increased abundance of the 5P reads at a given position indicates an increased ribosome pausing and slowed ribosome movement at that position on the mRNA. Relatively slower ribosome dynamics are observed in the WT (Black) compared to the mutant mH69 (Red) cells. **B. Metagene analysis displays the abundance of 5P reads relative to the stop codon in ORF**. The distribution of 5P reads relative to the stop codon in ORF. The significantly accumulated 5P reads, positioned 17 nt upstream of the stop codon, indicate the ribosome pause when a stop codone is encountered in the A site during translation termination. After this peak, the 3 nt periodicity associated with the ribosome movement on mRNAs diminishes as the translation process comes to an end. Notably, the peak indicating translation termination in the ribosome is higher compared to that for initiation in the ribosome. Similar to what is seen for translation initiation, slower ribosome dynamics are observed during termination in the WT (Black) compared to the mutant mH69 (Red). **C Ribosome pausing at the stop codons during translation termination**. Metagene analysis of ribosome pauses at the stop codons UAA, UGA and UAG. 5P read counts at each codon position are normalized to RPM (Reads Per Million), and data over the window spanning from - 30 to +2 are shown. The WT and mH69 are shown in black and red, respectively. The WT exhibits similar slower ribosome dynamics compared to mH69 at all three stop codons.

Compared to the subtle change observed in initiation, the difference in the ribosome dynamics is more pronounced at the stop codon during translation termination between mH69 and WT. In 5P-Seq experiments, observable ribosomal pauses during the termination step give rise to a clear peak positioned 17 nucleotides upstream of the stop codon, where the stop codon is positioned in the ribosomal A site (**Figure 3B**). Here, the WT ribosome clearly shows a greatly elevated levels of accumulated 5P intermediates at the -17nt position compared to the mutant mH69, suggesting faster ribosome dynamics in mH69 in termination. Consistent with this observation, earlier studies showed that defects in H69, including nucleotide mutations or missing rRNA modifications in this region, cause termination defects such as stop-codon read-through (18, 20). It is worth noting that the changes in ribosome dynamics in termination observed here are not specifically related to the identity of the stop codon encountered. Instead, our data show that all three stop codons, UAA, UGA and UAG, show similar defects in the mutant ribosome at the metagene level (**Figure 3C**). Taken together, these results suggest loss of Ψ2258 and Ψ2260 of H69 results in faster translation initiation and termination.

### 3. Translation over a subset of codons is particularly sensitive to the missing Ψ2258 and Ψ2260 of H69

Next, we asked whether there were any changes in ribosome dynamics when translating the 61 sense codons. To answer this question, we studied ribosome footprints in sense codon decoding by comparing the normalized 5P reads between WT and mH69 cells. Several observations follow: First, at the metagene level, we observe limited changes in ribosome footprints between mH69 and WT at the amino acid level at -17nt (A site), -14nt (P site) and -11nt (E site) (**Supplemental Figure S2**). Furthermore, when we analyzed reads over tripeptides, including consecutive charged amino acids as well as proline-rich peptides, we did not observe any statistically significant changes to the ribosome dynamics when Ψ2258 and Ψ2260 were absent in H69 (**Supplemental Figure S3)**.

Secondly, at the codon level, we identified changes in mH69 ribosome dynamics when decoding a subset of GC-rich codons during translation elongation (**Figure 4A**). Obvious translation pauses were found when mH69 ribosomes decode the arginine codons CGA and CGG in the A site. Consistent with our observations related to amino acids that were described above, these changes are codon-specific, rather than amino acid-specific, as decoding of the arginine codon CGT in the ribosome does not seem to be affected when Ψ2258 and Ψ2260 are lacking.

**Figure 4.**
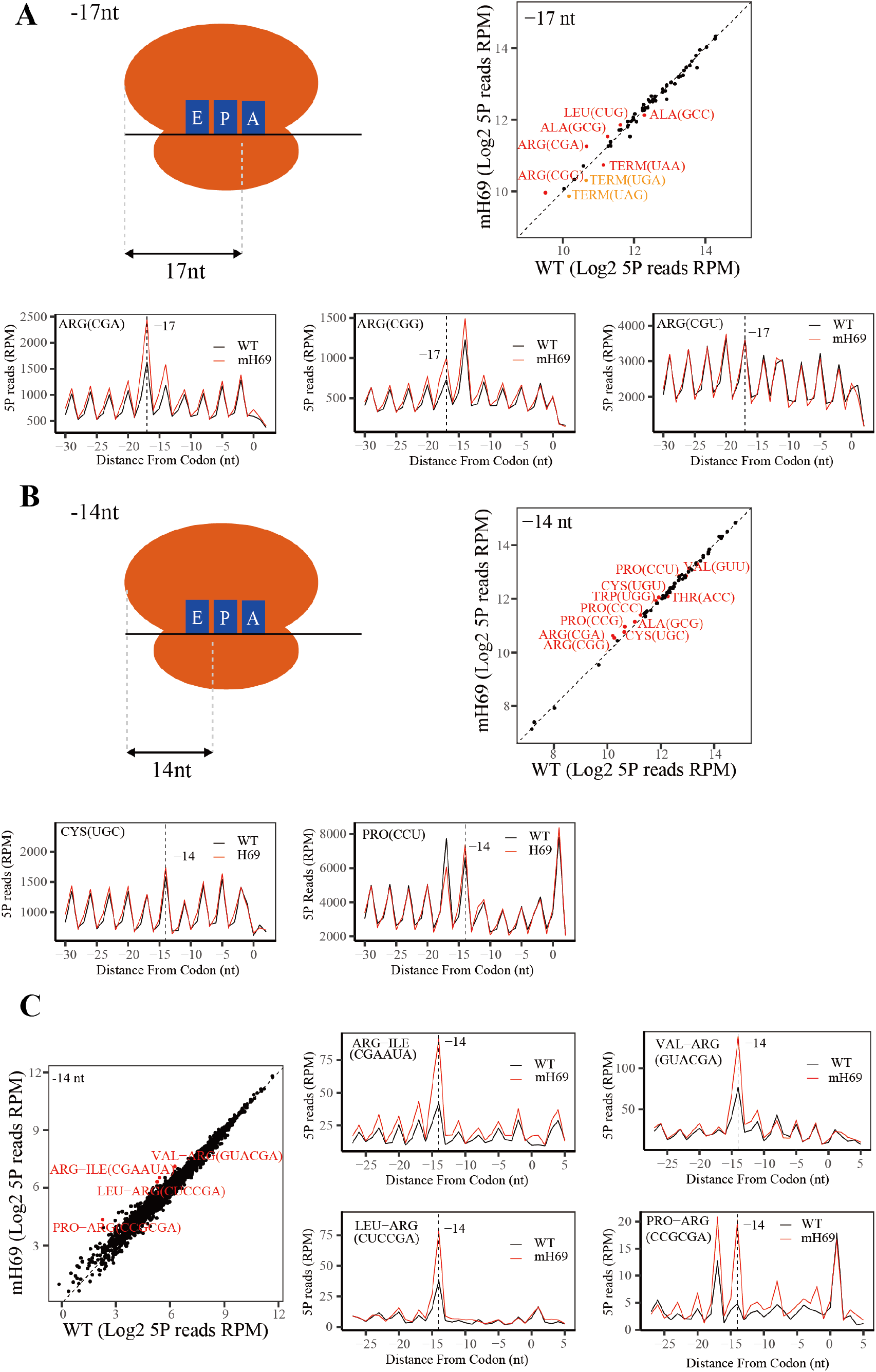
Loss of Ψ2258 and Ψ2260 in H69 alters ribosome dynamics during translation elongation. **A and B: Codon-specific changes of ribosome dynamics upon missing Ψ2258 and Ψ2260 of H69.** ***<u>A</u>*** *<u>and</u>* ***<u>B</u>*** *<u>Up Panel</u>*: Left: Schematic representation indicating the relative distance of the 5’ end of 5P reads to the ribosomal A-site (A) and P-site (B). Right: Scatter plots depicting differential ribosome pausing at specific codons at ribosome A-site (A) and P-site (B). The 5P read counts are normalized and converted to log2-RPM values. Codons showing statistically significant differences are marked in red (adjusted p-value < 0.01). Additionally, changes of 5P read counts at stop codons UGA and UAG with adjusted P-values < 0.05, are shown in orange. ***<u>A</u>*** *<u>and</u>* ***<u>B</u>*** *<u>Bottom Panel:</u>* Increased translation pauses at Arg codons CGA and CGG (**A**) and Cys (UGC) and Pro (CCU) (**B**) in the absence of Ψ2258 and Ψ2260 of H69. **C Di-codon-specific changes of ribosome dynamics when Ψ2258 and Ψ2260 are absent in H69.** Left: Scatter plots showing differential ribosome pauses at consecutive di-codons occupying ribosome A- and P-sites. Di-codons that show statistically significant changes (with adjusted p-value < 0.01) are shown in red. Right: Differential ribosome pauses of di-codons at the metagene level. The total number of reads was normalized to facilitate data analysis and comparison, and data were analyzed as described in Methods. The peaks at 17 nt, 14 nt, and 11 nt represent indicated codons at the ribosomal A-, P-, and E-sites, respectively. Positions of ribosome pauses at -17nt and -14nt are indicated with dashed lines. The normalized RPMs for each codon between the WT (black lines) and mH69 (red lines) cells were compared. The metagene (−30 to +2 window) shows the number of 5P intermediates from the translation of indicated codons and their ribosomal positions in mH69 (red lines) and WT (black lines) cells. The adjusted p value was calculated using the Benjamin-Hochberg method in R package (see Methods,). The adjusted p < 0.01 was used as the criterion for a significantly regulated difference in the data.

Thirdly, the presence of Ψ2258 and Ψ2260 in H69 is important for translation dynamics over many GC-rich codons residing in the ribosomal P-site. As shown in **Figure 4B**, a greater number of GC-rich codons at the 14nt position show differences (with statistical significance) in the normalized 5P reads between mH69 and WT cells, though the magnitude of those differences is generally smaller compared to those observed at the 17nt position (ie, ribosome A site).

Fourthly, consistent with these observations, but perhaps not surprisingly, GC-rich di-codons, particularly those containing the rare codon Arg (CGA), such as Arg-Ile (CGA-AUA), Val-Arg (GUA-CGA) and Leu-Arg (CUC-CGA), show obviously increased translation stallings/pauses when they are encountered in the ribosomal A and P sites simultaneously (**Figure 4C**). Notably, consecutive rare codons demonstrated a marked stalling in the ribosome when Ψ2258 and Ψ2260 of H69 are missing. As shown in the bottom panel of **Figure 4C**, mH69 mutant ribosomes with rare di-codon CCG-CGA (Pro-Arg) in the P and A sites show a significantly increased level of stalling.

Finally, we do not find any changes to the ribosome dynamics in mH69 in the ribosomal E site (−11nt position) (**Supplemental Figure S3)**, nor when Arg or Pro-rich amino acids have been incorporated into the nascent peptides in the ribosomal exit tunnel (data not shown).

Of note, since the data show different codons for the same amino acid are differentially affected, the observed ribosome pauses likely resulted from insufficient tRNA availability or altered ribosome-tRNA interactions, rather than the availability of amino acids. As an example, the Arg(CGA) codon, which is the rarest codon in Saccharomyces cerevisiae, decreases the efficiency of translation due to the wobble decoding by tRNA^Arg (ICG)^ (24, 25). Coexpression of an exact match tRNA mutant, tRNA^Arg(UCG)^, was reported to completely suppress the inhibitory effect rendered by this rare codon (24). Since H69 contacts tRNAs in both A and P sites in the ribosome, it is conceivable that the presence of the two pseudouridines, Ψ2258 and Ψ2260, contribute to the dynamics of the H69 RNA stem-loop, and help to maintain functional interactions with the tRNAs as they move within the ribosome. The optimality of this interaction is likely to be more critical for those limited tRNAs that decode rare codons in an organism.

### 4. Loss of Ψ2258 and Ψ2260 gives rise to a decrease in the IRES-dependent translation

To further characterize changes in the ribosome dynamics upon loss of these two conserved central pseudouridines, we studied the internal ribosome entry site (IRES)-dependent translation activity by the mutant ribosome. IRES-dependent translation initiation by the Dicistroviridae family IRESs, such as that employed by Cricket paralysis virus (CrPV), has been studied extensively (26, 27). The viral CrPV-IRES recruits the ribosome directly to the mRNA in the absence of protein initiation factors. The mechanism of IRES-dependent translation initiation employed by CrPV viral RNA is particularly sensitive to ribosome dynamics. In addition to the binding of the IRES to the ribosome, a “pseudotranslocation” event allows CrPV to move its start codon from the ribosomal A to P site, which is important for CrPV to reach an elongation-competent translation state (27, 28). Similar to tRNA translocation in the ribosome, pseudotranslocation occurs with the assistance from elongation factor eEF2 and GTP hydrolysis and involves structural rearrangement in the ribosome (27, 28). Conceivably, ribosome dynamics play a critical role in IRES translocation (29, 30).

We performed a cell-free translation assay using a reporter mRNA harboring a CrPV-IRES element in the 5’UTR of an in-frame luciferase gene. snoRNAs play diverse cellular functions beyond ribosome biogenesis, such as alternative splicing and microRNA-like functions (31-35). To rule out other such mechanisms that could alter luciferase activities through indirect effects resulting from snR191 deletion, we purified WT and mutant mH69 ribosomes and combined them with wildtype supernatants to generate a reconstituted cell-free translation cocktail (**Figure 5A**). This way, only the direct effect on mRNA translation conferred by Ψ2258 and Ψ2260 modifications to the ribosome can be studied. As shown in **Figure 5B**, CrPV-IRES-mediated translation activity was much reduced in the mutant ribosome. In contrast, translation activity of a capped luciferase mRNA shows insignificant changes in the mutant ribosome (**Figure 5C)**. Consistent with these results, it is reported that hypopseudourinated ribosomes are defective in translating mRNAs containing IRES elements (36-38).

**Figure 5.**
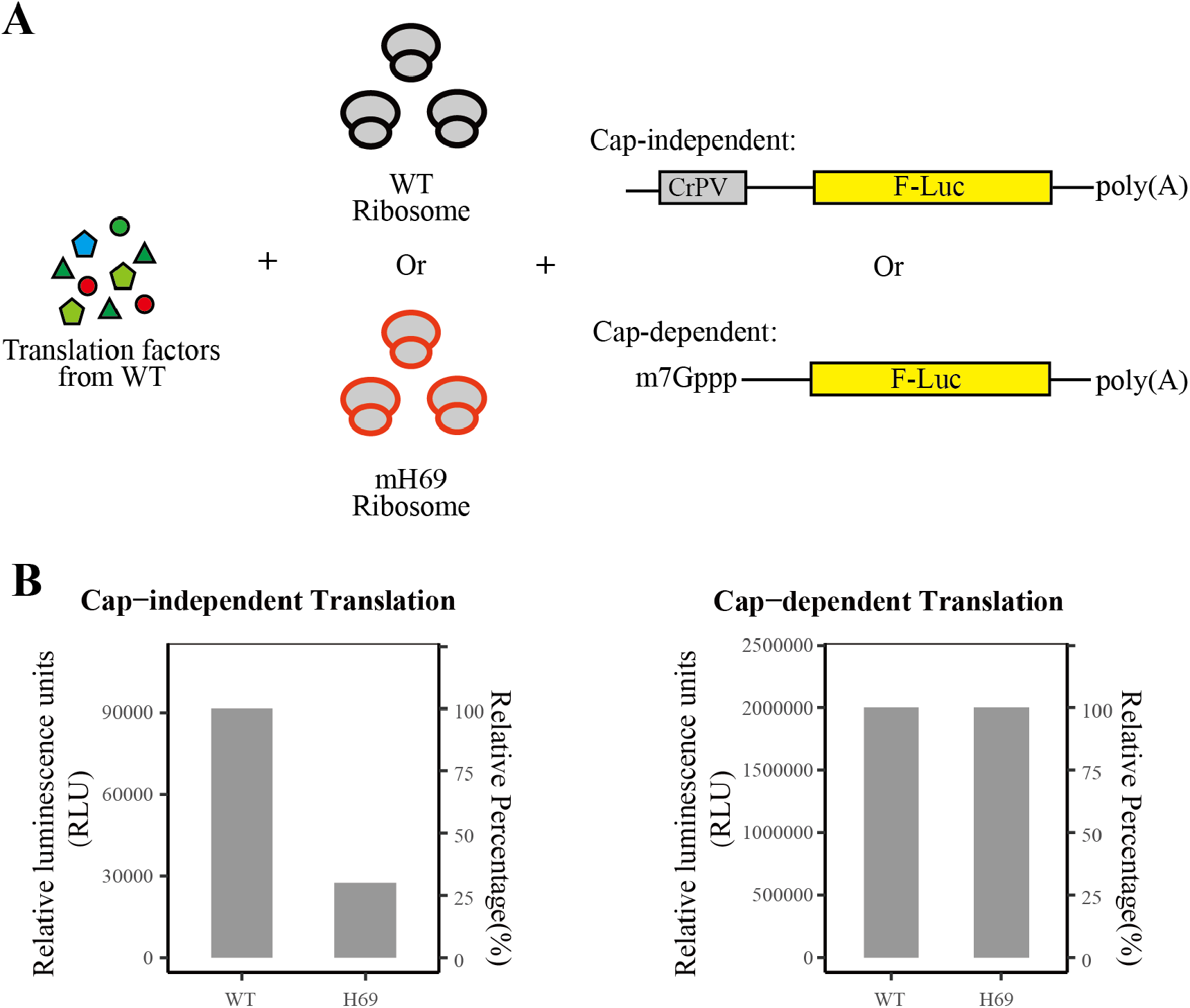
Loss of Ψ2258 and Ψ2260 in H69 impairs IRES-mediated translation. **A. Schematics of the cell-free *in vitro* translation assay using reconstituted WT and mH69 ribosomes with WT S100**. WT and mH69 ribosomes were purified and reconstituted with the wildtype S100 (WT cell lysates devoid of their native ribosomes). Cap-dependent and CrPV(IRES)-dependent translation activities of the WT and the mH69 ribosomes were measured using luciferase mRNAs containing a 5’ Cap or CrPV(IRES) in the 5’UTR, respectively. **B. Loss of Ψ2258 and Ψ2260 in H69 impairs IRES-dependent translation**. CrPV(IRES)-dependent (Left) and Cap-dependent (Right) translation activities of the WT and mH69 ribosomes were obtained. Our data show missing Ψ2258 and Ψ2260 in H69 impairs CrPV(IRES)-dependent translation but leaves the cap-dependent translation initiation activity largely unchanged. All of the measured activities were normalized to the amount of luciferase mRNAs and the total proteins present in the reconstituted system.

## DISCUSSIONS

The ribosome is a highly conserved ribonucleoprotein machine that carries out codon-dependent protein synthesis in all cells. The fundamental mechanism of protein synthesis in the ribosome heavily depends on ribosomal RNAs rather than ribosomal proteins. Taking the three conserved functional centers of the ribosome as an example, the peptidyl transfer center, the decoding center and the GTPase-associated center are all formed by rRNAs only, with proteins decorating in the vicinity. rRNAs are also the key players that mediate subunit interactions, also known as “bridges” in the ribosome. Furthermore, functionally important regions of rRNA are often subject to covenant modifications. Many of these modifications are conserved, suggesting their functional importance. In this study, we have investigated the functions of the highly conserved pseudouridines in H69 of the ribosome.

H69 is a universally conserved RNA helix in the large ribosomal 28S rRNA (23S rRNA in bacteria). Deletion of H69 results in a dominant lethal phenotype (39), indicating its essential cellular function. The loop region of H69 contacts the minor groove of the stem of h44 in the 18S rRNA (16S rRNA in bacteria) that is immediately adjacent to the ribosomal decoding center. This interaction forms the inter-subunit bridge B2a which is conserved across all kingdoms of life and is also present in organellar ribosomes, such as those within mitochondria and chloroplasts. The integrity of the B2a bridge is shown to be critical for ribosome function, as nucleotide mutations or rRNA modifications in H69 cause defects in subunit association (10, 40, 41) and impair translation accuracy (10, 11).

RNA nucleotides in H69 are heavily modified. In S. cerevisiae, there is one methylated ribose (Am2256) and four pseudouridines (Ψ2258, Ψ2260, Ψ2264 and Ψ2266) in this region (**Figure 1A**), all of which are conserved from yeast to humans. In particular, the presence of pseudouridines at positions 2258 and 2260 (Ψ1915 and Ψ1917 in H69 of the 23S rRNA in E. coli (42)) is a highly conserved feature of the ribosomes from bacteria to humans, suggesting their functional significance to the cell. Although the absence of these modifications can be tolerated under standard laboratory growth conditions, the mutant cell cannot compete with the wild-type (21), showing that pseudouridylation at these positions confers the fitness to the cell.

Functions of modified nucleotides are traditionally studied using methods such as cell growth assays and standard reporter-based assays. Here, using a new genomic method called 5P-Seq which probes *in vivo* ribosome dynamics on mRNAs, we obtain the translation profiles of yeast cells missing these two conserved ribosomal pseudouridines (Ψ2258 and Ψ2260 in H69). Our results show changes in the ribosome dynamics upon loss of these modifications in the center of the ribosome. Specifically, we observe faster translation initiation and termination at the metagene level, and the mutant ribosomes are more prone to frameshifting. Although ribosomes lacking these two modifications exhibit relatively faster movement, our data also indicate that they stall more frequently than those with the modifications when translating GC-rich codons, especially the rare codons.

Loss of the rigidifying effect that is conferred by pseudouridines in H69 allows for greater conformation flexibility of H69, which likely accounts for the faster ribosome dynamics during mRNA translation. Pseudouridines stabilize local secondary or tertiary structure (43, 44), and the resulting fine-tuning of RNA structure may facilitate the RNA function (45). The synergistic effects of clusters of modifications seen in the functional regions of the RNA are also consistent with this notion. The chemical differences between uridine and pseudouridine largely account for these effects. Although having a similar base-pairing capacity as uridine, pseudouridine possesses an “extra” hydrogen bond donor at its N1 position (**Figure 1B**). This donor can form a hydrogen bond with a water (H2O) molecule that can in turn form a hydrogen bond with other molecules (46, 47). Thus formation of pseudouridine can provide a subtle, but potentially important rigidifying influence that enhances favorable base stacking (48-50) and stabilizes higher-order interactions via H2O mediated hydrogen bonding via (? through) the N1 amino proton (51-55) in RNAs.

Defects in pseudouridine synthase (Cbf5 in yeast and dyskerin DKC1 in mammalian cells) diminish the number of pseudouridines in cellular RNAs. Ribosomes isolated from these cells are under-pseudouridylated and showed a decreased affinity for their tRNA ligands and increased programmed ribosome frameshifts (15, 16). Decreased IRES-mediated translation was observed in the hypopseudouridylated (15, 16, 37, 38) and hypomethylated ribosomes (13, 14). However, it is striking to see that even missing two conserved pseudouridines, Ψ2258 and Ψ2260, in H69, results in a drastic decrease in the IRES-mediated translation. The central location of the two pseudouridines likely accounts for this observation. Consistent with this, H69, due to its central location, is essential for the stability of the ribosome. In the absence of tRNA ligands, the deletion of H69 completely abolishes the association of the two ribosomal subunits, even at high magnesium concentrations, and allows for *in vitro* recycling of the two ribosomal subunits in the absence of ribosome recycling factor (39). Therefore, it is perhaps not surprising that even subtle changes in H69 result in substantial alterations of ribosome stability and dynamics in translation.

Earlier studies showed that ΔsnR191strain cannot compete with the WT cell (21). The observed loss of fitness in mutant cells can be explained, at least partially, by compromised protein homeostasis due to changes in ribosome dynamics. Deviations from the optimum cellular protein homeostasis, although seemingly negligible under standard laboratory conditions, may lead to detrimental consequences *in vivo* and manifest as severe disease phenotypes in patients.

Furthermore, impaired IRES-mediated translation can also contribute to the loss of fitness of mutant cells. Initially discovered in viruses (56, 57), IRES-like elements/structures are also found in the 5’ untranslated region (5’UTR) of eukaryotic mRNA transcripts, many of which encode functionally important proteins involved in critical cellular processes. IRES-like elements are functionally important for enabling alternative cap-independent translation (58), which is critically important during cellular processes such as developmental patterning, cell cycle progression, and stress conditions when the cap-dependent translation pathway is inhibited. In line with this explanation, compromised translational regulation of IRES-containing oncogenes and tumor-suppressor genes by hypopseudouridylated ribosomes leads to tumorigenesis (37) and defects in oncogene-induced senescence (OIS) (38).

Because of the conserved nature of Ψ2258 and Ψ2260 in H69 of the ribosome, it is possible that the defects in ribosome dynamics upon loss of these pseudouridines may also be conserved from bacteria to humans. Furthermore, the manifestation of these defects, such as defects in maintaining the correct open reading frames, achieving optimal tRNA interactions IRES-mediated translation, are also likely to be conserved.

On the other hand, the cognate snoRNA, snR191, which guides the formation of these two modifications, is highly conserved (21). Although deletion of snR191 only showed minor growth defects (21), snoRNA deletion can affect ribosome biogenesis and mature ribosome formation (59). It remains to see whether snR191 deletion affects other cellular pathways beyond translation. Further investigations are also required to understand specific interactions that pseudouridines have with molecular signatures of tRNAs, mRNAs, and translation factors, and to provide mechanistic insights into enhanced translation by these covalent modifications.

## METHODS

### Yeast Strains

*S. cerevisiae* strains YS602 (MATα ade2-101 trp1-Δ101 ura3-52 leu2-3, 112 his3Δ200) and mH69 (ΔsnR191) are generous gifts from M.J. Fournier at UMass Amherst. We also deleted snR191 from the WT YS602 strain following the published method in the lab (18).

### In vitro translation assay

The cricket paralysis virus IRES was cloned upstream of the firefly luciferase reporter gene, with aT7 promoter upstream and polyA tail downstream. Linearized templates were obtained by PCR using a T7 sequence (TAATACGACTCACTATAGG) as the forward primer and a poly(dT) with 60 d(T) as the reverse primer. The PCR product was purified by phenol chloroform extraction and RNA was synthesized using the HiScribe T7 Quick High Yield RNA Synthesis Kit (NEB). 1 ug of starting linear DNA template was used and transcription was performed for 2hrs at 37°C. The product RNA was precipitated by LiCl precipitation for 30min at -20°C, and the pellet obtained by centrifugation for 15min and resuspended in Tris-HCl, pH 6.5. The RNA was flash-frozen and stored at -80°C until use. Agarose bleach gel electrophoresis was performed to evaluate the integrity of RNA. For capped luciferase mRNA, commercial CleanCap Fluc mRNA was used (Trilink).

Yeast cells were harvested at an OD_600_ value of 0.6 and were lysed in buffer containing 30mM Hepes, pH 7.4; 100mM KOAc; 2mM Mg(OAc)_2_; 2mM DTT and protease inhibitor (complete protease inhibitor cocktail tablet, EDTA free, Roche). Ribosomes were pelleted for 2.5 hours at 176k x g. mH69 and wildtype ribosomal pellets were resuspended in wildtype supernatant depleted of ribosomes. For *in vitro* translational assays, S100 were treated with 9000U/ml micrococcal nuclease and 2.5mM CaCl_2_ for 7min at 26ºC (60). The mRNAs were denatured at 95ºC for 30 sec and snap-cooled before being added to the translation reaction. 1µg of either capped or IRES-containing mRNA was mixed with treated S100 lysate with reconstituted ribosomes of equal protein concentration, as measured by OD280. Reactions were conducted a buffer containing 22 mM Hepes-KOH (pH 7.4), 120 mM KOAc, 2 mM Mg(OAc)_2_, 0.75 mM ATP, 0.1 mM GTP, 25 mM creatine phosphate (Invitrogen), 0.04 mM of each of the twenty amino acids, 1.67 mM 1,4-dithiothreitol (DTT), 5 μg creatine kinase (Invitrogen) and 20U RNase inhibitor (NEB). The translation reactions were incubated in 384 well plates at 30°C for 1hr. The luciferase activity was measured with ONE-Glo Luciferase reagent (Promega) with an Analyst HT plate reader. The measurements were normalized by calculating the total protein amount in each well using Bradford reagent (Biorad).

### 5P-Seq

Co-translational mRNA degradation intermediates were captured from mH69 and WT cells using 5PSeq (22). Yeast cells were grown in 50ml YEPD and harvested at an OD600 value of 0.6. Harvested cells were resuspended in a lysis buffer containing 50 mM Tris-HCl (pH 8.1), 130mM NaCl, 5mM EDTA, and 5% SDS. Cells were lysed using glass beads and total mRNA was extracted using phenol-chloroform. 50ng of total mRNA was used as the starting material to perform 5PSeq. Poly(A)-containing mRNA was enriched using 100µl oligo(dT) beads (Dynabeads oligo (dT)25, Life Technologies) for 5P samples while randomly generated fragments of total RNA were prepared as controls. Both 5P and control RNAs were fragmented at 80°C for 5min. Following reverse transcription and second strand generation, overnight incubation at 4ºC with M-280 streptavidin beads (Life Technologies) was performed instead of 30min at room temperature, as suggested for maximum capture of the cDNA. The final DNA adapter ligation reaction was carried out for 2hrs at 16°C. Following PCR, each sample was purified separately with HighPrep beads (Magbio) as suggested and their quality was checked using Agilent 2100 Bioanalyzer. A Qubit 2.0 Fluorometer (Life Technologies) was used to determine the concentration of each library. Libraries were pooled in equal molar ratios and fragments were enriched between 300-500nt for the library. The final libraries were sequenced using Illumina NovaSeq 6000.

### Data processing

Adapters were trimmed from sequenced reads using cutadapt v1.18 (61) and barcodes were split using fastx barcode splitter in the Fastx-toolkit. Unique molecular identifiers (UMI) were extracted and deduplicated using UMI tools v1.01 (62). Reads were aligned to the Saccharomyces cerevisiae.R64-1-1.107 reference genome (Ensemble) by STAR v2.7.3a(63). BedGraph profiles and RNA content were separated by bedtools v2.29.2 (64) and SAMtools v1.10.2 (65), respectively. Read counts were quantified for each gene using htseq-count v 0.11.3(66) and reads were visualized on genome tracks using Integrative Genomics Viewer(IGV) (67). The fivepseq 1.0.0 (68)was employed to analysis 5’ends positions. Differentially expressed genes or codons were conducted in R v.4.2.3. Counts were normalized to reads per million (RPM)), and subsequently, the DEseq2 was utilized to identify differentially expressed genes or codons (69). The resultant p-values were adjusted using the Benjamini-Hochberg method to evaluate the significance of the findings. Gene ontology (GO) enrichment analysis was performed using clusterProfiler v.4.0.5 package in R (70). A false discovery rate (FDR) threshold of 0.05 was established as the cut-off for significance.

## ACKNOWLEDGEMENTS

We thank Roy J. Carver Biotechnology Center at the University of Illinois at Urbana-Champaign for sequencing, and members in the Jin laboratory for helpful discussions. We thank M. Chowdhury for pioneering yeast in vitro translation assay in the lab and C.W Hawk for helpful comments on the manuscript.

