## Supplemental Figures for "Conserved Pseudouridines in Helix 69 of the Ribosome are Important for Ribosome Dynamics in Translation"

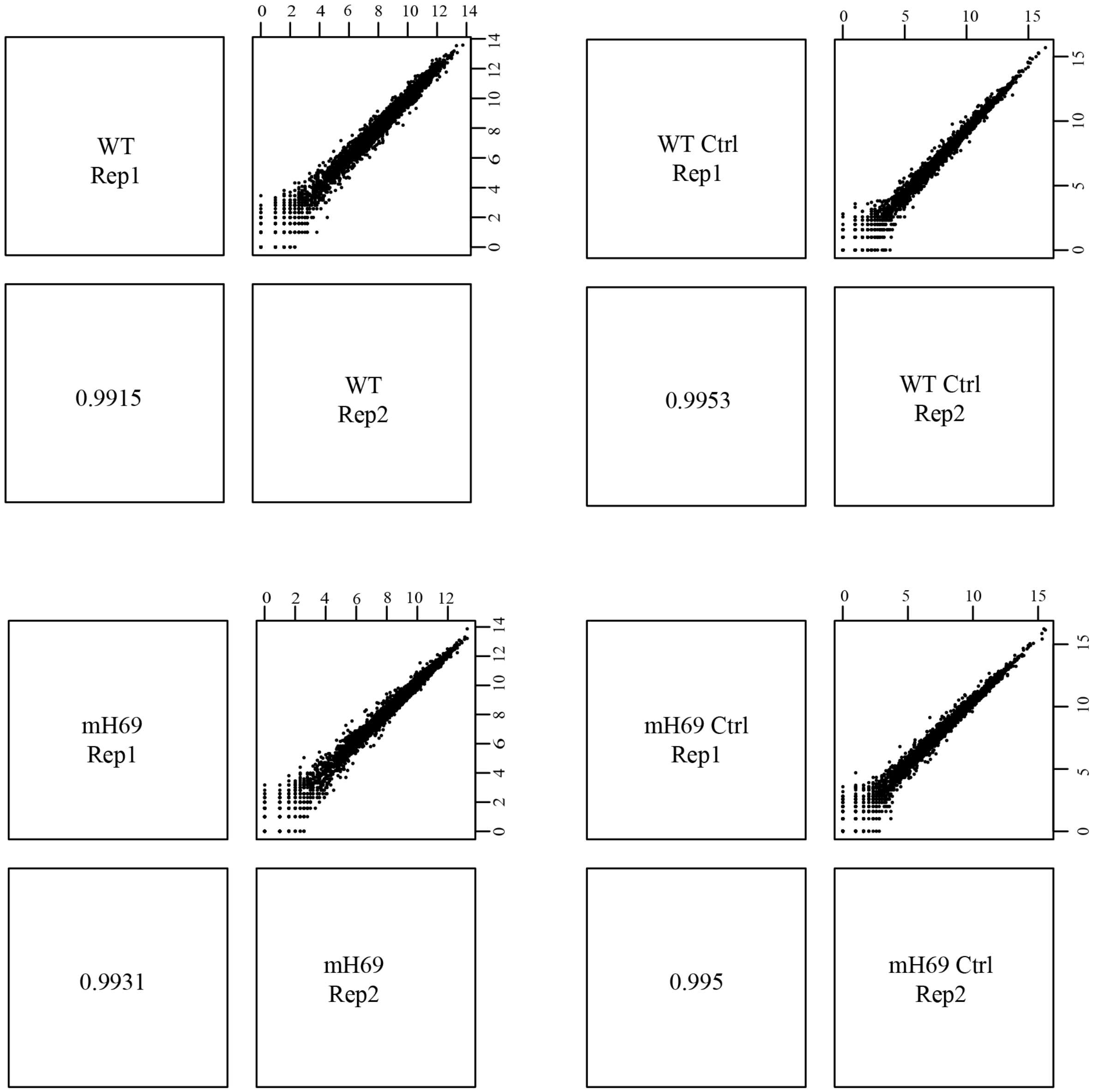


**Supplemental Figure S1: Data quality and reproducibility of the 5P-Seq Experiments**

Scatter plots illustrate normalized 5P reads (on the log2 RPM (Reads Per Million) scale) to evaluate the reproducibility of 5P-Seq experiments for WT and mH69. Corresponding controls are also included. Results for the two biological replicates are compared and Pearson correlation coefficients for each comparison are shown.

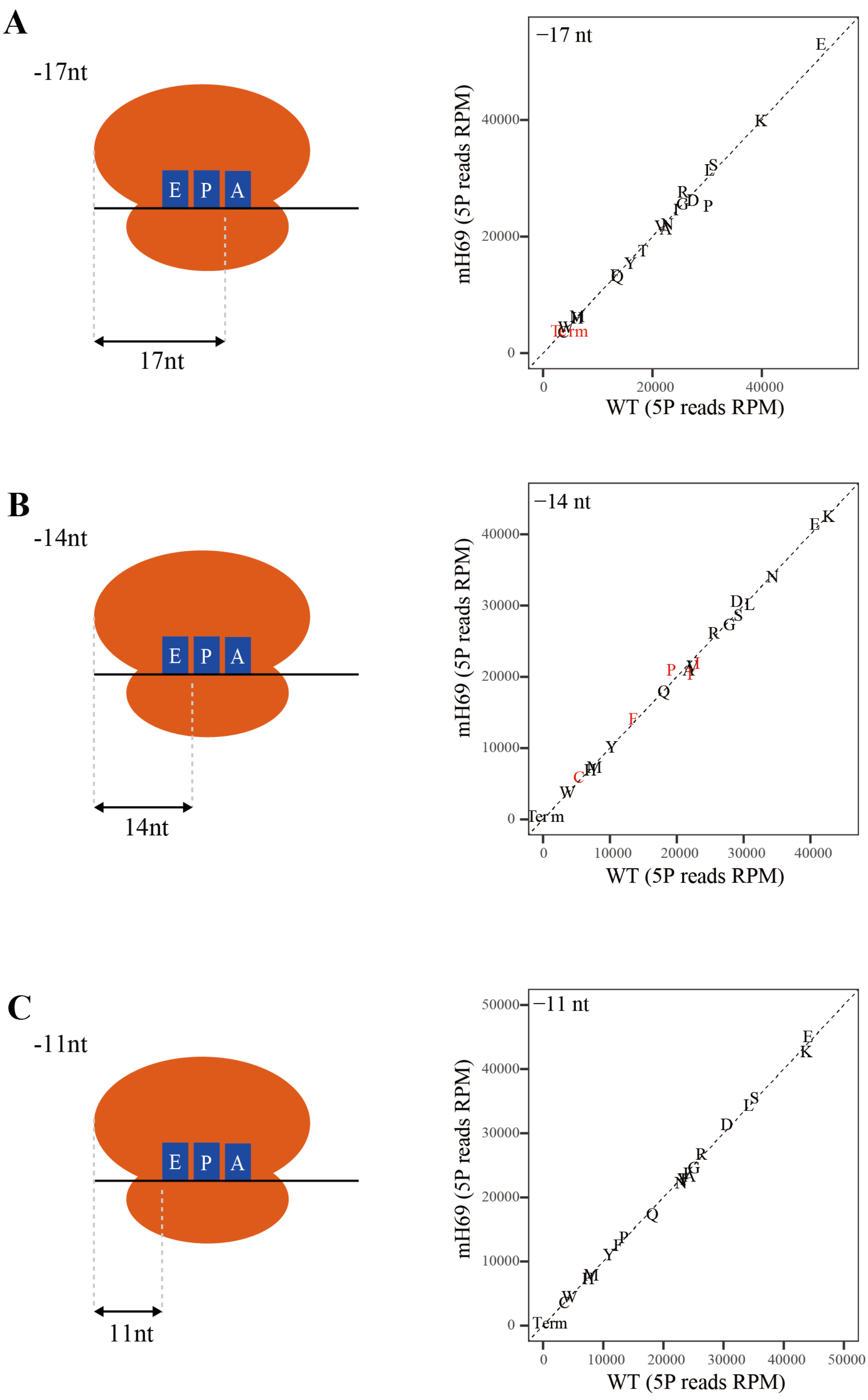
A
-17nt
E
P
A
17nt
−17 nt
mH69 (5P reads RPM)
WT (5P reads RPM)
0
20000
40000
Term
B
-14nt
E
P
A
14nt
−14 nt
mH69 (5P reads RPM)
WT (5P reads RPM)
0
10000
20000
30000
40000
Term
C
-11nt
E
P
A
11nt
−11 nt
mH69 (5P reads RPM)
WT (5P reads RPM)
0
10000
20000
30000
40000
50000
Term

**Supplemental Figure S2: Loss of Ψ2258 and Ψ2260 of H69 confer little changes to translating ribosome dynamics at the amino acid level.**

**A, B and C**: Left: Schematics depicting the corresponding distances between the ribosomal sites occupied by codons (& tRNAs) and their sequence reads captured by 5P-Seq. the 5’ end of 5P reads mapped to 17nt, 14nt and 11nt upstream indicate protected ribosome footprints when translating codons occupying at the ribosomal A (A), P (B) and E (C) sites, respectively. Right: The differential ribosomal pausing for all amino acids at -17 nt (A-site codon), -14nt (P-site codon) and -11nt (E-site codon). Statistically significant changes with adjusted p-value $< 0.01$ are shown in red.

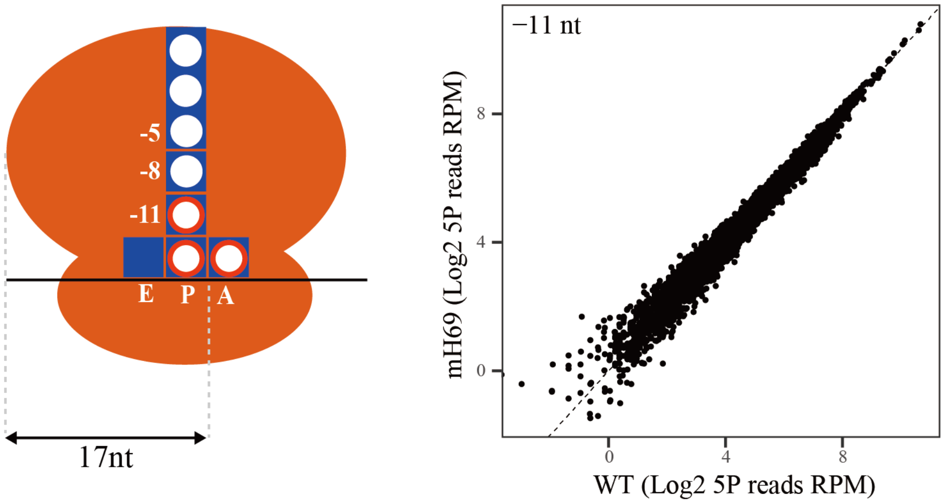


**Supplemental Figure S3: Impact of tripeptide motifs in mH69 translation**

Left: Schematic depicting the number of consecutive peptides that occupy the ribosome during translation. Amino acids are represented as white circles. Positions of each amino acid relative to the 5' end of the 5P reads are labeled accordingly. Tripeptide motifs of interest are outlined in red.

Right: Differential ribosome pausing across all tripeptide motifs in the genome in WT and mH69 cells. The WT 5P reads in log2-RPM (Reads Per Million) are plotted on the x-axis, and those for mH69 are plotted in the same scale on the y-axis. Data analysis reveals that none of the tripeptide motifs show statistically significant changes (adjusted p-value $< 0.01$) in mH69 lacking Ψ2258 and Ψ2260 of H69.
